# SuperMetal: A Generative AI Framework for Rapid and Precise Metal Ion Location Prediction in Proteins

**DOI:** 10.1101/2025.03.21.644685

**Authors:** Xiaobo Lin, Zhaoqian Su, Yunchao (Lance) Liu, Jingxian Liu, Xiaohan Kuang, Peter T. Cummings, Jesse Spencer-Smith, Jens Meiler

## Abstract

Metal ions, as abundant and vital cofactors in numerous proteins, are crucial for enzymatic activities and protein interactions. Given their pivotal role and catalytic efficiency, accurately and efficiently identifying metal-binding sites is fundamental to elucidating their biological functions and has significant implications for protein engineering and drug discovery. To address this challenge, we present SuperMetal, a generative AI framework that leverages a score-based diffusion model coupled with a confidence model to predict metal-binding sites in proteins with high precision and efficiency. Using zinc ions as an example, SuperMetal outperforms existing state-of-the-art models, achieving a precision of 94 % and coverage of 90 %, with zinc ions localization within 0.52 ± 0.55 Å of experimentally determined positions, thus marking a substantial advance in metal-binding site prediction. Furthermore, SuperMetal demonstrates rapid prediction capabilities (under 10 seconds for proteins with **∼** 2000 residues) and remains minimally affected by increases in protein size. Notably, SuperMetal does not require prior knowledge of the number of metal ions—unlike AlphaFold 3, which depends on this information. Additionally, SuperMetal can be readily adapted to other metal ions or repurposed as a probe framework to identify other types of binding sites, such as protein-binding pockets.

## 1 Introduction

The Protein Data Bank (PDB) contains nearly 200,000 structures, and approximately one-third of these proteins contain metal ions [1]. Many proteins require the binding of one or more metal ions to perform their functions. Zinc, a vital biologically active metal, is particularly noteworthy as it binds to approximately 10% of all human proteins [2]. These proteins rely on zinc for their biological function, structural stability, or regulation of activities. For example, zinc is indispensable for the activity of more than 300 enzymes, spanning all six enzyme classes [3]. The binding of zinc also stabilizes the folded conformations of protein domains, ensuring proper function [4]. Additionally, zinc ions play a crucial role in cell-cell communication, cell signaling, proliferation, survival, and DNA repair by modulating zinc-binding proteins [5]. In mammals, zinc homeostasis is primarily maintained by zinc transporter proteins, and zinc deficiency may lead to various health issues, such as smell and taste disorders, immune system disorders, developmental delays, and impaired immune function [6].

Given the importance and unique functionality of zinc in proteins, accurately identifying zinc-binding sites is crucial. However, direct determination of these sites through wet experiments is often costly and time-consuming due to technical difficulties, requiring expensive instruments, complex procedures, and elaborate labor. In contrast, computational approaches for discovering zinc-binding sites can significantly reduce the cost and time associated with corresponding wet experiments. Consequently, many computational methods have been developed to predict zinc-binding sites, which can be broadly categorized into four groups based on the attributes they consider: (1) template-based methods that leverage structural or sequence homologs and binding templates, e.g., MIB and MIB2 [7, 8]; (2) sequence-based methods that rely on amino acid sequence, e.g., M-Ionic [9]; (3) structure-based methods that use structural geometry and chemical features to identify metal-binding sites, e.g., Metal1D, Metal3D, and BioMetAll [10, 11]; and (4) physics-based methods, such as quantum mechanical/molecular mechanical (QM/MM) simulations [12, 13]. However, template-based methods are limited by the availability of existing templates or patterns and may struggle to accurately predict novel binding sites, while sequence-based methods lack the ability to offer atomic-level insights or detailed descriptions of protein-metal interactions. Physically modeling the dynamics of metal ion binding sites is also challenging, especially for sites located in the protein interior [14, 15]. Moreover, physical methods like molecular dynamics simulations struggle to find appropriate force fields for transition metals to reproduce correct coordination distances [14, 16, 11], while QM/MM simulations are too computationally expensive to be practical for typical protein design tasks. [12, 17, 18].

Current state of the art predictors for metal location is Metal3D [11], a structure-based method that employs 3D Convolutional Neural Networks (CNNs) to predict the positions of metal ions, such as zinc. Metal3D operates by taking a protein structure along with a specified set of residues as input, voxelizing the environment around each residue, and then predicting the per-residue metal density. Trained on experimental zinc sites, Metal3D generates a probability density map for metal ions, offering high precision with predictions within sub-angstrom precision. Despite its success, Metal3D faces challenges similar to other 3D CNN models [19, 20, 21, 22], such as the need for fine grid spacing (voxelization). The computational cost for these voxel-based models increases cubically with the resolution of the input, making scaling up difficult [23]. Moreover, these CNN-based models are sensitive to the orientation of the input structure, requiring data augmentation to increase the number of training samples and reduce the risk of overfitting [11, 24]. Although Metal3D has demonstrated superior accuracy compared to other methods, issues like rotational invariance remain, highlighting the need for further advancements in metal ion prediction models [25, 26]. In recent years, diffusion models have emerged as a powerful generative AI technology with applications spanning natural language processing, image synthesis, and bioinformatics [27, 28, 29]. These diffusion models have significantly advanced fields such as computational protein design [30], drug and small-molecule development [31], protein–ligand interaction modeling [28, 29], cryo-electron microscopy data enhancement [32], and single-cell data analysis [33, 34]. In contrast, other generative approaches—such as variational auto-encoders (VAEs)—often require strict architectural constraints to maintain a tractable normalizing constant or must rely on surrogate objectives for approximate maximum likelihood training [35]. Meanwhile, generative adversarial networks (GANs) typically use adversarial training, which is notoriously unstable and prone to mode collapse [36]. By directly modeling the gradient of the log probability density, score-based diffusion models avoid many of these pitfalls, offering a more stable way to represent probability distributions.

In this paper, we introduce SuperMetal, a novel generative AI approach that combines a score-based diffusion model with equivariant graph neural networks to accurately predict zinc ion positions within protein structures. Instead of directly approximating the probability distribution of zinc ions, our model estimates the gradient of this distribution and then uses it to generate zinc positions from a normal distribution. These generated positions are then refined through a confidence model and clustered, resulting in a precise prediction of both the number and exact locations of zinc ions for a given protein structure. SuperMetal surpasses existing methods, achieving state-of-the-art results in the coverage of experimental zinc ions and the precision of predicted positions. This approach offers significant potential for applications in structural studies, binding site predictions, multi-body docking, and metalloprotein engineering.

## 2 Results and Discussion

In this section, we present the evaluation and comparative results of SuperMetal against the state-of-the-art Metal3D in predicting zinc ion positions within protein structures. Both methods utilize the same test dataset to ensure a fair comparison. During the inference step of SuperMetal, 100 metal ions are sampled at random positions across the system and denoised via reverse diffusion over their translational degrees of freedom. A confidence model and clustering mechanism are subsequently applied to determine the final zinc ion binding sites for different proteins.

### 2.1 Overview of SuperMetal

The SuperMetal pipeline consists of three primary stages. The process begins with the preprocessing of protein structures into heterogeneous geometric graphs. These graphs consider node and edge features for protein residues, protein atoms, and metal ions, while also accounting for diffusion time at different timesteps [37, 38]. These geometric graphs are then fed into a score-based diffusion model to sample potential zinc ion positions. Following the sampling, an equivariant graph neural network is utilized to predict confidence scores for each candidate metal position. Positions with confidence scores below a specified threshold are subsequently filtered out. In the final stage, a clustering algorithm [39] is employed to calculate the centroids of neighboring metal positions, resulting in the placement of one metal ion per cluster.

The detailed methodology for each of these steps is provided in Section 4 and the supplemental information, with a visual representation of the entire pipeline illustrated in Fig. 1. The key contributions of SuperMetal include:

**Fig. 1.**
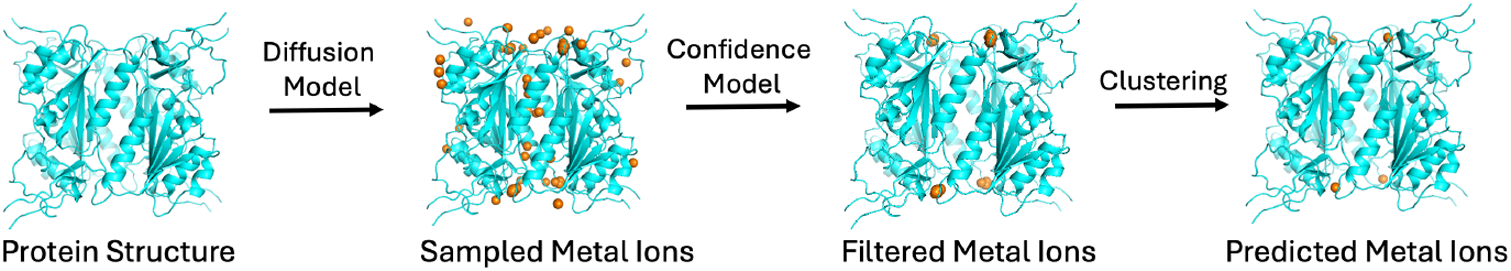
Workflow of SuperMetal. The orange spheres represent the sampled Zn ions. The protein, shown in blue, is from 2J9R in the RCSB Protein Data Bank.

- A score-based diffusion model that processes geometric graphs of protein structures, enabling the sampling of metal positions within proteins.
- An equivariant graph neural network that accurately evaluates each sampled point and filters out low-confidence positions.
- Postprocessing operations designed to optimize prediction accuracy by clustering the predicted positions.

### 2.2 Comparison between SuperMetal and Metal3D

In the field of ligand-binding site prediction and water site prediction, 3D-CNN–based deep-learning models have demonstrated superiority over knowledge-based, sequence-based, physics-based, and traditional machine-learning methods [19, 20, 40]. Metal3D [11] has been reported to achieve state-of-the-art performance in predicting zinc ion binding locations. The tool was benchmarked against other existing methods, including Metal1D [11], BioMetAll [10], MIB [8], and MIB2 [41]. Metal3D operates by taking a protein structure along with a specified set of residues as input, voxelizing the environment around each residue, and then predicting the per-residue metal density. The predicted densities for each residue, within a 16 × 16 × 16 Å^3^ volume, are averaged to compute the overall zinc density across the protein. Clustering of voxels exceeding a defined probability threshold is then performed based on the global probability density. For each resulting cluster, a weighted average of the voxel positions is computed, using the probability at each point as the weight, ultimately placing one metal ion per cluster [11].

In contrast, SuperMetal employs a fundamentally different approach to predict metal ion binding locations. We conceptualize metal binding as a generative modeling problem—learning a distribution over metal translations given multiple known metal ion locations and the target protein structure. To address this, we developed SuperMetal, a diffusion generative model (DGM) that operates in the translational space of metal locations for metal binding prediction. This model defines a diffusion process over the translational degrees of freedom, specifically the positions of various metal ions relative to the protein. SuperMetal generates metal positions by running a learned reverse diffusion process, which iteratively transforms an initial, noisy prior distribution over metal positions into a refined model distribution (Fig. S2). Conceptually, this can be seen as the progressive refinement of random initial positions through successive updates of their translations. Additionally, we train a confidence model to estimate the likelihood of the metal positions generated by the DGM, allowing us to select the most probable samples. The final predicted metal locations are determined by calculating the cluster centers. The last two-step process significantly enhances prediction accuracy while maintaining a minimal impact on computational runtime.

It is important to note that SuperMetal operates in a continuous space, eliminating the need to voxelize protein environments, as is necessary with Metal3D. In Metal3D, the voxel grid size and resolution can significantly influence prediction accuracy and computational cost. Metal3D also requires rotational augmentation of the residue environment during voxelization to enhance data generalization and the training of machine learning models. In contrast, SuperMetal leverages the full 3D structure, allowing the scoring model to reason about physical interactions using an SE(3)-equivariant framework without the need for data augmentation [42], which enhances its ability to generalize to previously unseen complexes.

To assess the model’s performance, we calculated the precision versus coverage curve for SuperMetal and Metal3D (see Section 4.5 for details). Predictive models typically exhibit a trade-off between precision and coverage [43], which is consistent with our observations in Fig. 2. The curve in this figure is generated using different probability thresholds for both SuperMetal and Metal3D. SuperMetal demonstrates higher precision across a wider range of coverage compared to Metal3D. For example, when Metal3D achieves 100% precision, its coverage is around 30%, whereas SuperMetal reaches approximately 70% coverage at the same level of precision—more than double the coverage of Metal3D. Similarly, at 77% coverage, SuperMetal maintains near 100% precision, while Metal3D’s precision drops to around 93%. Moreover, at 88% coverage, Metal3D’s precision is ∼84%, whereas SuperMetal achieves ∼95%, marking a significant improvement. These results clearly indicate that SuperMetal outperforms Metal3D, providing higher precision even at greater coverage. This demonstrates SuperMetal’s ability to maintain high precision while significantly expanding the scope of its predictions.

**Fig. 2.**
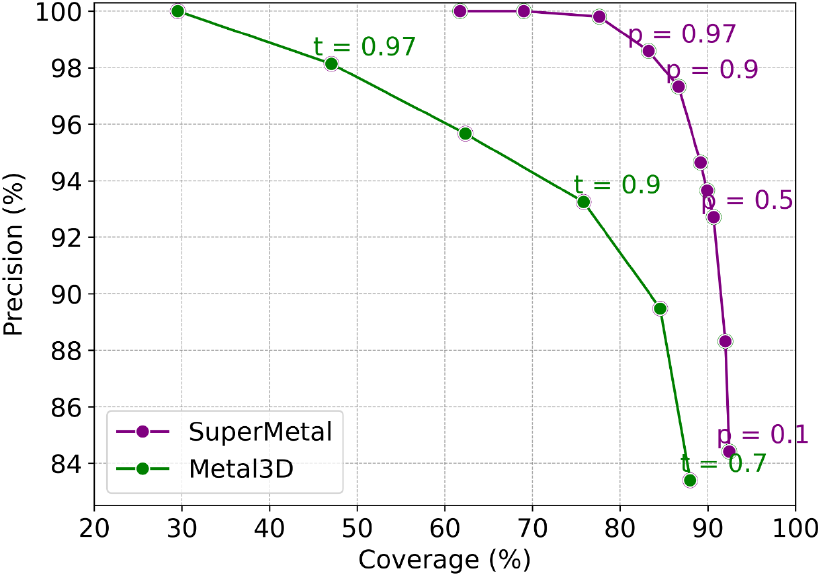
Precision Versus Coverage for SuperMetal and Metal3D at Different Probability Cutoffs. The graph compares SuperMetal (purple line) and Metal3D (green line), illustrating how precision varies with coverage at different cutoff values. The labels indicate the probability cutoffs, *p* for the confidence model in SuperMetal and *t* for Metal3D.

In addition to precision and coverage in metal site prediction, it is crucial to assess the spatial precision of these predictions. To evaluate this, we measured the mean absolute deviation (MAD) between the experimentally determined zinc locations and their corresponding correctly predicted positions (true positives), as shown in Fig. 3. At a probability threshold of *p* = 0.1, SuperMetal achieves a MAD of 0.61 ± 0.66 Å, which improves to 0.44 ± 0.58 Å as the threshold increases to *p* = 0.9. This trend demonstrates that higher probability cutoffs lead to greater spatial precision, with the median MAD decreasing from 0.37 Å at *p* = 0.1 to 0.23 Å at *p* = 0.999. The relatively small difference between these two median values suggests that even low-confidence predictions are spatially accurate within the protein structure. In contrast, Metal3D exhibits a clearly increasing median MAD, rising from 0.36 Å at *t* = 0.7 to 0.87 Å at *t* = 0.99, indicating a greater deviation from ground-truth positions as the probability cutoff increases. Additionally, the spread of MAD values in SuperMetal decreases with higher probability cutoffs, opposite to the trend observed in Metal3D, where the spread increases.

**Fig. 3.**
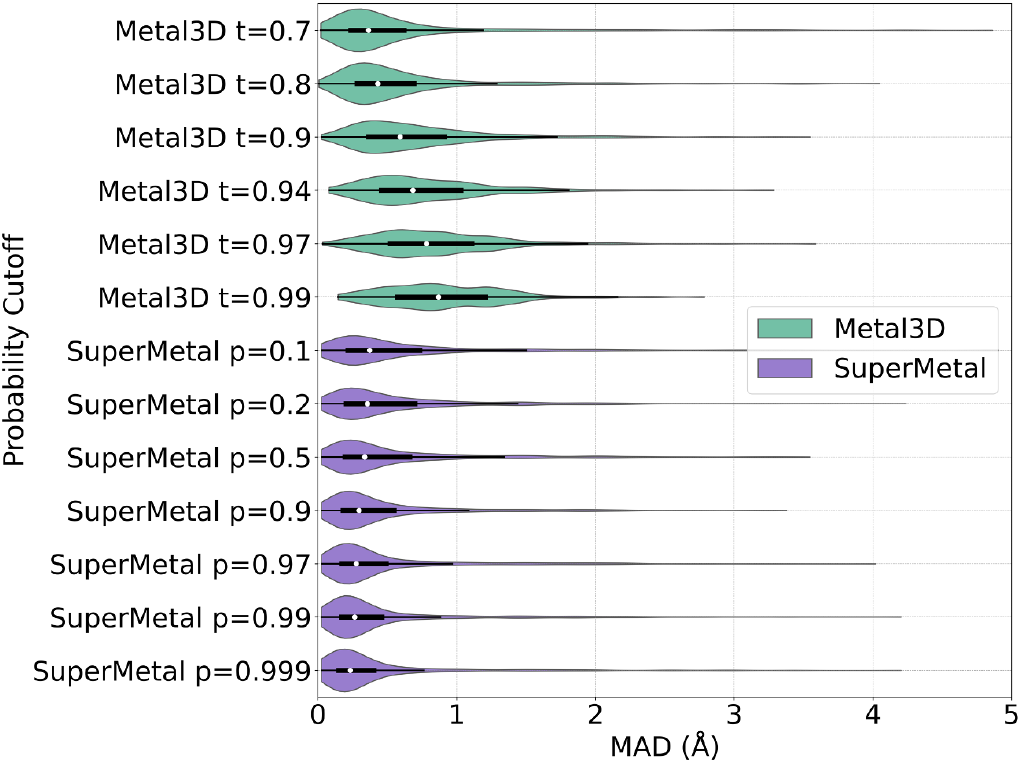
Distribution of Mean Absolute Deviation (MAD) for SuperMetal and Metal3D at Increasing Probability Cutoffs. This violin plot illustrates the MAD for metal ion location predictions by SuperMetal and Metal3D, with cutoffs ranging from lower to higher values for each model. SuperMetal is shown in purple and Metal3D in green. The plot employs kernel density estimation to display the data distribution, with the white circle indicating the median, the black box defining the first and third quartiles, and whiskers extending up to 1.5 times the interquartile range to capture the spread of typical data values.

These observations from the precision vs. coverage and MAD analyses suggest that SuperMetal not only demonstrates strong spatial precision across varying probability cutoffs, but also reveals that its improved precision in metal site prediction is closely tied to enhanced spatial precision, as indicated by a smaller MAD. In other words, as SuperMetal’s confidence in its predictions increases, so does its precision in pinpointing the exact locations of metal-binding sites within protein structures. This level of precision is particularly critical in applications where accurate metal ion positioning is essential, further highlighting the robustness and reliability of SuperMetal’s predictive capabilities.

SuperMetal not only outperforms Metal3D in terms of prediction accuracy but also demonstrates a significant advantage in processing speed. The computational runtime comparison as a function of protein size (number of residues) for SuperMetal and Metal3D is shown in Fig. 4. For consistent comparison, both models were executed using a single thread on one CPU core, with the same GPU. We observed that Metal3D’s runtime tends to increase exponentially as the protein size grows, whereas SuperMetal maintains consistently low runtimes (under 10 seconds), even for larger proteins. For example, when the protein size approaches 2000 residues, Metal3D requires approximately 500 seconds, which is around 60 times longer than SuperMetal.

**Fig. 4.**
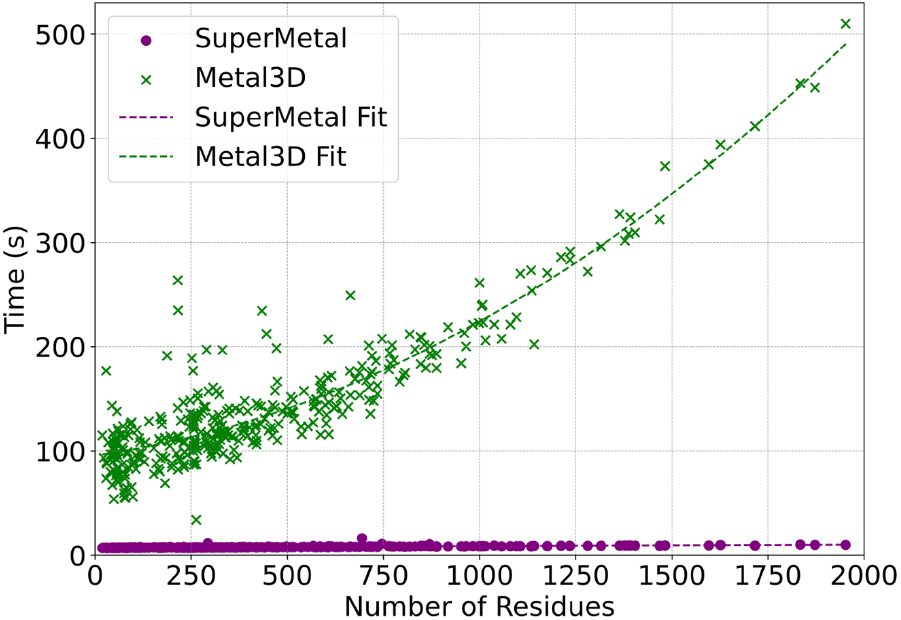
Computational Time Analysis for SuperMetal and Metal3D Across Protein Sizes. This scatter plot compares the computational time required by SuperMetal and Metal3D to predict metal-binding sites against the number of protein residues. Polynomial regression curves (purple and green dashed lines) are only used to clarify the trends.

This impressive performance improvement in SuperMetal can be attributed to the multi-scale approach that SuperMetal employs. Unlike Metal3D, SuperMetal efficiently manages computational complexity through a hierarchical interaction system. For instance, when metal ions are distant from protein residues, only coarse-grained interactions are considered; when metal ions are closer to the residues, the atomic structure of the residues is taken into account. This approach ensures that SuperMetal avoids constructing overly large and computationally expensive graphs, particularly between the metal and protein or within the protein itself. By limiting the scope of interaction calculations to only those that are most relevant and proximal, SuperMetal significantly enhances its computational efficiency. In contrast, Metal3D employs a different approach, utilizing voxelization and grid averaging for proteins. While this method is effective in certain contexts, it results in computation times that scale more directly with the number of residues. Consequently, as protein size increases, Metal3D experiences a much more rapid rise in runtime, leading to significantly longer processing times compared to SuperMetal, particularly as protein complexity increases.

### 2.3 Case study

The recently published AlphaFold 3 [44] has garnered significant attention due to its ability to predict interactions within joint structures of complexes, such as protein-ligand and protein-ion interactions. In this case study, we compare our model, SuperMetal, against AlphaFold 3 and Metal3D, focusing on zinc-binding site prediction. For the evaluation, we selected two proteins with distinct zinc-binding sites: 5IN2, which represents the crystal structure of the extracellular Cu/Zn superoxide dismutase enzyme from Onchocerca volvulus [45], and 6BTP, the crystal structure of bone morphogenetic protein 1 (BMP1) complexed with a hydroxamate inhibitor [46].

One challenge we encountered with AlphaFold 3 is the need to specify the exact number of zinc ions for binding site prediction. In real-world scenarios, the number of ion-binding sites is often unknown, making this requirement impractical. To facilitate comparisons, we specify the number of input zinc ions for AlphaFold 3, using one, two, and six zinc ions for each protein (from left to right, as shown in Fig. 5). Another significant limitation of AlphaFold 3 is that its source code is not publicly available. This lack of access restricts users from modifying or customizing the tool to meet specific needs or integrate it with other workflows. Moreover, AlphaFold 3’s server only accepts sequence inputs, not direct PDB structures, which can be problematic when the protein structure is already known, as using the structure would be preferable in such cases. However, for the two protein structures used in the case study, we fortunately found that the structures predicted by AlphaFold 3 from sequences were quite similar to the experimentally determined PDB structures. This allowed us to use AlphaFold 3’s results for comparison with Metal3D and our model (see Fig. 5).

**Fig. 5.**
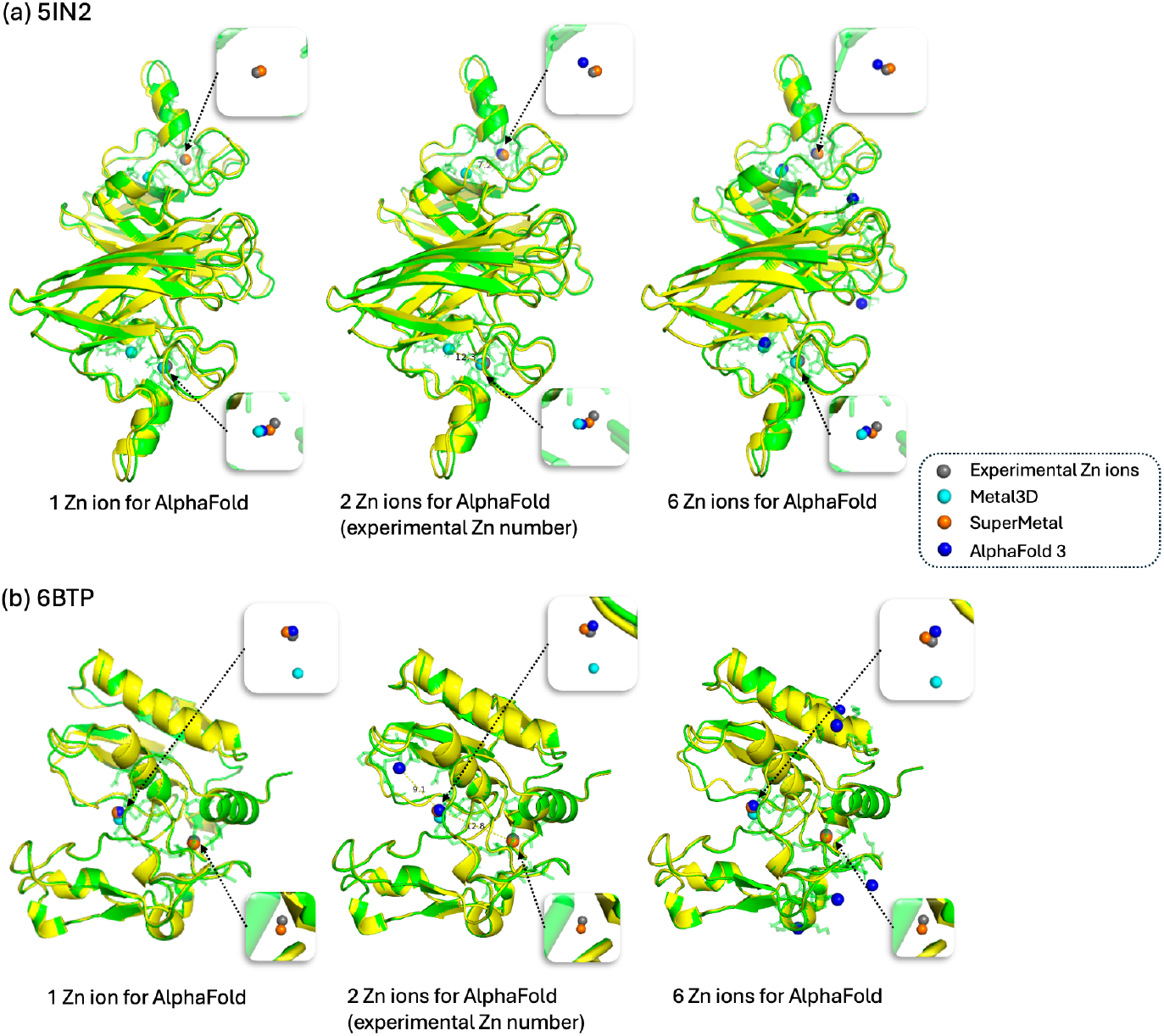
Comparative visualization of Zn ion binding site predictions for the proteins 5IN2 (1) and 6BTP (2). Zn ions are color-coded as follows: grey for experimentally determined Zn ions, cyan for Metal3D predictions, orange for SuperMetal predictions, and blue for AlphaFold 3 predictions. The protein structure is shown in green for the same input PDB file used in Metal3D and SuperMetal, while it is shown in yellow for AlphaFold 3. The transparent green region around the Zn ions highlights the protein’s atomic structure within a 5 Å radius of the metal ions. From left to right, the figure shows varying numbers of Zn ions specified in the AlphaFold 3 input, ranging from 1 Zn ion, to 2, and finally to 6 Zn ions.

**Fig. 6.**
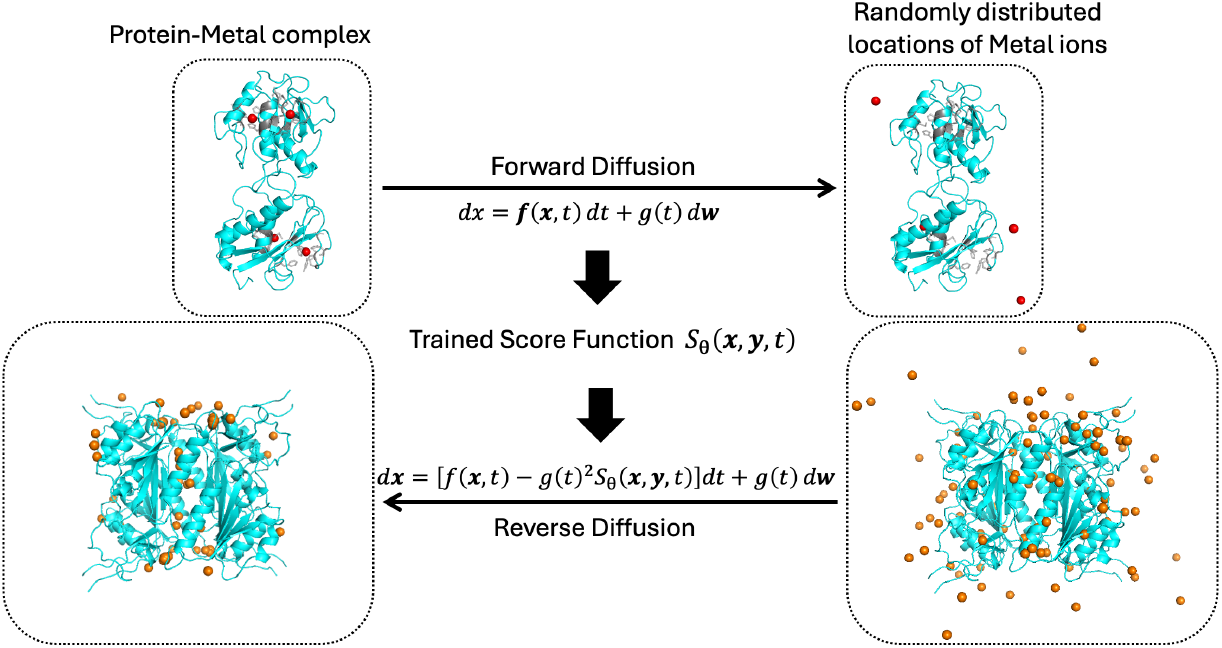
Fundamental Theory of the Score-based Generative Diffusion Model for Metal Ions in Proteins. The forward continuous-time stochastic differential equation (SDE) transitions the true locations of metal ions (top left) to random locations (top right). The score at each intermediate time step is predicted by a deep learning neural network, enabling the reverse process of the SDE. The grey part of the protein (top) represents the atomic structure of the protein surrounding the metal ions at their original positions.

In the case of 5IN2, when we specified the experimentally determined number of zinc ions (i.e., 2), AlphaFold 3 accurately predicted the zinc-binding locations with 100% precision and 100% coverage (Fig. 5a, Table 1). However, this approach relies on prior knowledge of the exact number of zinc ions, which is not always feasible in real-world applications. For instance, when AlphaFold 3 was provided with only one zinc ion, the precision remained high (100%), but the coverage dropped significantly to 50%. Conversely, inputting six zinc ions into AlphaFold 3 led to multiple incorrect zinc-binding predictions (Fig. 5, Table 1). However, even when the correct number of zinc ions was specified, AlphaFold 3’s predictions were not always reliable: in the case of 6BTP, despite inputting the experimental determined number of zinc ions, both precision and coverage reach only 50%. In contrast, SuperMetal consistently achieved 100% precision and coverage in this scenario, demonstrating its robustness in handling such predictions.

**Table 1.** Comparison of precision and coverage in predicting zinc ion binding sites using Metal3D, SuperMetal, and AlphaFold 3.

| Model | 5IN2 |  | 6BTP |  |
| --- | --- | --- | --- | --- |
|  | Precision (%) | Coverage (%) | Precision (%) | Coverage (%) |
| Metal3D | 33 | 50 | 100 | 50 |
| SuperMetal | 100 | 100 | 100 | 100 |
| AlphaFold 3 (1 Zn ion) | 100 | 50 | 100 | 50 |
| AlphaFold 3 (2 Zn ions) | 100 | 100 | 50 | 50 |
| AlphaFold 3 (6 Zn ions) | 33 | 100 | 17 | 50 |

Our results also show that SuperMetal outperformed Metal3D in both precision and coverage across the two case studies, reinforcing the findings from the previous section.

## 3 Conclusions

In this work, we introduced SuperMetal, a novel generative AI pipeline for predicting metal ion positions within protein structures. Our method leverages a score-based diffusion model to sample potential metal positions, followed by an equivariant graph neural network as a confidence model to evaluate and refine these predictions. The effectiveness of SuperMetal was tested against the state-of-the-art Metal3D model using a comprehensive dataset of zinc-binding sites.

SuperMetal demonstrated superior performance across all key metrics, including precision, coverage, and mean absolute deviation (MAD). Notably, SuperMetal consistently outperformed Metal3D by maintaining high precision while achieving significantly higher coverage, even at stricter thresholds. For instance, at a precision of ∼100%, SuperMetal achieved nearly double the coverage compared to Metal3D, demonstrating its ability to identify a larger set of correct metal positions without compromising precision. Additionally, SuperMetal showed improved spatial precision, as reflected by consistently lower MAD values across all probability cutoffs. Beyond its predictive precision, SuperMetal also excels in scalability, maintaining low inference times across varying protein sizes. Unlike Metal3D, which exhibits an exponential increase in runtime with increasing protein size, SuperMetal efficiently handles large proteins using a multi-scale approach that reduces computational complexity. Furthermore, SuperMetal predicts metal-binding locations without requiring prior knowledge of the number of metal ions, a limitation present in AlphaFold 3, which relies on this information for its predictions. This flexibility adds further utility to the SuperMetal framework, distinguishing it from existing methods.

These results underscore the robustness and versatility of SuperMetal, establishing it as a valuable tool for accurately predicting metal ion positions in protein structures. The ability to reliably pinpoint these positions carries significant implications for understanding metal-dependent biological processes, protein stability, and enzyme catalysis.

Despite the promising results achieved by SuperMetal, several limitations remain. First, our metalbinding site predictions currently rely on data from the ZincBind dataset, which focuses on zinc-binding proteins. Second, our approach does not explicitly incorporate additional structural elements such as RNA, small molecules, or water. Addressing these issues is critical for ensuring broader applicability across diverse biological scenarios. Future work could involve expanding the dataset to include a wider range of metal ions and structural properties. Additionally, linking holo (bound) complexes with their corresponding apo (unbound) and predicted structures may further strengthen the model’s generalizability and support more realistic inference scenarios. In principle, our framework can readily be adapted to complex biological contexts. For example, we recently extended our generative AI framework and data curation strategies to predict water-binding sites in proteins, as well as protein–ligand and protein–protein complexes, demonstrating state-of-the-art performance in those domains [47].

Lastly, although our training data included proteins up to ∼3000 residues—covering the majority of biologically relevant metalloproteins, given that those exceeding 1500 residues are relatively rare [48]— SuperMetal supports structures of any size, provided sufficient memory is available to construct the required all-atom and coarse-grained graphs. For exceptionally large proteins or resource-constrained environments, users can segment the protein into smaller sections, run predictions piecewise, and then merge the results.

## 4 Methods

### 4.1 Data Set and Preprocessing

For this study, we utilize the ZincBind database [49], a high-quality, non-redundant collection of zinc-binding sites extracted from the RCSB Protein Data Bank (PDB) [50]. The database comprises 19,154 unique sites across 19,103 PDB files. To eliminate redundancy, ZincBind clusters zinc binding sites based on the structural similarity of zinc-binding chains and by comparing the amino acid sequences and types within the binding site. Furthermore, the database includes only zinc sites satisfying physiologically relevant criteria (e.g., at least two liganding residues and three liganding atoms), and accounts for symmetrical units in protein structures, thereby preventing the misclassification of essential catalytic or structural zinc ions as merely surface zinc ions [51]. From the 19,154 unique sites, we extracted 10,253 PDB files, each containing one or more of these sites. Structures with more than 3000 residues were excluded from the dataset. Additionally, all PDB structures were cleaned by removing any exogenous ligands, retaining only the zinc ions. In cases where multiple models were available for a given structure, such as those with alternative residue conformations, only the first model was used. We then randomly selected 1,000 structures for validation. Separately, our test set comprises 350 structures, including those from the original Metal3D test set and additional structures randomly sampled from our non-redundant dataset. To ensure a fair comparison, none of these 350 test structures contain binding sites similar to those in the training sets for both SuperMetal and Metal3D. Testing was conducted under identical computational conditions using one CPU and one GPU (Nvidia A100 40 GB).

### 4.2 Diffusion Model

Each data point in the diffusion model represents the three-dimensional coordinates of a metal ion’s position within a protein structure [27, 52, 53]. The goal of the generative model is to approximate the probability distribution *p*(**x** | **y**), where **x** refers to the metal positions and **y** denotes the protein structure [28]. Approximating *p*(**x** | **y**) presents two main challenges.

The first challenge is that directly computing the probability distribution is intractable, as it requires normalizing the distribution across the entire space of possible positions. To circumvent this, instead of estimating *p*(**x** | **y**) directly, the diffusion model is utilized to estimate its gradient, ∇_**x**_ log *p*(**x** | **y**), commonly known as the score function *S*_*θ*_(**x**) [54, 55]. The score function *S*_*θ*_(**x**) is a vector field that describes the direction in which the metal ions should move to reach their favorable positions from a given point in 3D space.

The second challenge arises from the lack of sufficient training data in certain regions of the protein. To address this, the true data distribution is ‘evolved’ into a known distribution, typically a normal distribution [27, 56]. Through this process, the model diffuses data (i.e., metal ions) across the entire three-dimensional space conditioned on the protein structure, effectively filling in knowledge gaps and learning from a broader range of information across the protein structure. The entire architecture is depicted in the Supporting Information.

#### 4.2.1 Forward Diffusion and Training

This evolution is facilitated by the forward step of the diffusion process, which is governed by a forward stochastic differential equation (SDE), described as:

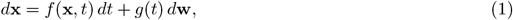

where **x** represents the positions of all metal ions, *t* denotes time, **w** refers to Gaussian noise or Brownian motion, *g*(*t*) is the diffusion coefficient, and *f* (**x**, *t*) is the drift coefficient. In our case, *f* (**x**, *t*) = 0, and *g*(*t*) is defined as 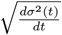. The variance *σ*^2^(*t*) evolves according to a hyperparameter of the model:

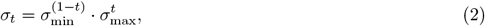

leading to the forward SDE:

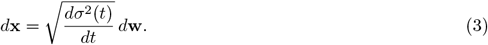

In other words, for a given protein-metal complex, as Gaussian noise **w**(*t*) is added and given the original distribution of metal ion positions at *t* = 0, denoted as **x**(0), the metal position at time *t*, **x**(*t*), can be numerically calculated through the equation above. Translation perturbation vectors, Δ**r**, also follow a Gaussian distribution with the mean *µ*(*t*) and variance *σ*^2^(*t*), which allows us to compute the gradient of the log probability of metal translations over the protein structure **y** using the following equation:

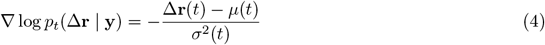

Meanwhile, the score function *S*_*θ*_(**x**) is generated by a neural network, where the inputs to the network are the locations of the metal ions, the protein structure, and the time *t*. The parameters of the neural network, *θ*, are optimized by minimizing the loss function *L*_*θ*_:

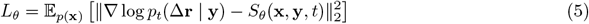

The expectation value 𝔼_*p*(**x**)_ is calculated by averaging the *L*_2_-norm between the true vector field ∇log *p*_*t*_(Δ**r** | **y**) and the predicted vector field *S*_*θ*_(**x, y**, *t*) across the metal distribution in the training data. We trained the model for 400 epochs employing this loss function.

#### 4.2.2 Reverse Diffusion and Sampling

In the forward diffusion process, given a protein structure, metal ions diffuse from their most favorable positions toward a Gaussian distribution in 3D space as noise is gradually added. During this phase, the score model learns the relationship between the favorable distribution at *t* = 0 and the perturbed distribution at time *t*.

Now, by reversing time, we use the well-trained score function *S*_*θ*_(**x**) to solve the reverse stochastic differential equation (SDE) and compute the favorable positions of metal ions from a random distribution:

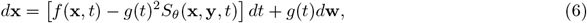

where *S*_*θ*_(**x**, *t*) is the learned score function from the training phase, and *f* (**x**, *t*) and *g*(*t*) represent the drift and diffusion terms from the forward SDE. For each protein structure, we perform inference by randomly initializing 100 candidate metal ion positions across the protein, ensuring that even sparsely represented or atypical coordination environments are sampled. These ions are guided by the score model *S*_*θ*_(**x**), ultimately reaching their most favorable positions within the protein structure.

### 4.3 Confidence Model for Filtering

The confidence model in SuperMetal is built on an SE(3)-equivariant convolutional network [42], adapted from the diffusion model architecture. Its primary objective is to classify metal ion positions sampled from the score model as either ‘good’ or ‘bad’ quality for filtering purposes. A “good” position is defined by having a mean absolute distance (MAD) below a certain threshold.

To train the confidence model, multiple samples are generated for each training complex using the trained diffusion model, and the MAD for each sampled metal ion position is computed. Labels are assigned by comparing these MAD values to the defined threshold, which in our experiments is set to 5 Å. The model is trained using cross-entropy loss to classify positions accordingly. The final layer of the confidence model employs a fully connected layer applied to the mean-pooled scalar representations from the last convolutional layer, producing an SE(3)-invariant confidence score. This score predicts the likelihood that a candidate metal ion position is favorable, ensuring that low-confidence positions can be filtered out. Accurate prediction of these confidence scores is essential for the post-processing step, as it allows for the precise distinction and placement of metal ions within the protein structure. The detailed architecture is provided in the Supporting Information.

### 4.4 Clustering Mechanism

The final step of the SuperMetal pipeline focuses on optimizing prediction precision through clustering of the predicted metal positions. After obtaining the sampled metal ion positions from the diffusion model, along with their confidence scores from the confidence model, a refinement process is applied. Positions with confidence scores below a defined threshold are filtered out, and the remaining high-confidence positions are clustered. A cluster is formed if it contains at least two metal positions within a 5 Å radius, using DBSCAN [57] algorithm implemented in scikit-learn [58]. For the DBSCAN algorithm, we set *ϵ* (the maximum distance between two points to be considered part of the same cluster) to 5 Å, consistent with the threshold for true positive metal predictions, and `min samples` (the minimum number of points required to form a cluster) to 2, as sampled metal locations are relatively sparse. For each cluster, the final metal ion position is determined by averaging the coordinates of all metal ions within the cluster, resulting in one metal ion per cluster.

### 4.5 Evaluation and Comparison

To evaluate the quality of predicted metal ion positions, we use precision, recall (coverage), and mean absolute deviation (MAD) as our primary metrics. These metrics provide a direct and intuitive measure of how accurately we identify metal-binding sites and how closely they match experimentally determined positions. In metal-binding site prediction, capturing all actual sites (i.e., maximizing recall) often takes precedence over penalizing false positives, which may reflect unobserved or transient sites.

Here, Precision is defined as the ratio of correctly predicted metal sites (true positives, TP) to the total number of predicted sites, including both correctly predicted sites and false positives (FP). Mathematically, precision is expressed as:

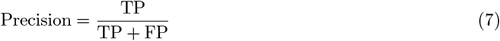

Recall, or coverage, measures the percentage of correctly predicted metal sites relative to the total number of true metal sites. This is represented as the ratio of true positives (TP) to the sum of true positives and false negatives (FN), where FN represents the true metal sites that were not detected:

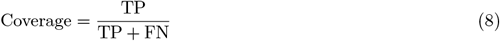

A metal site is considered correctly predicted (TP) if the predicted position falls within 5 Å of the experimentally determined site. False positives (FP) are predicted sites that do not meet this criterion, while false negatives (FN) are true metal sites that the model fails to predict.

To quantify the positional precision of the predictions, we calculate the mean absolute deviation (MAD). We chose MAD over Mean Squared Error (MSE) because MAD is more straightforward to interpret in the context of spatial distances and provides a clear measure of prediction accuracy in terms of physical distance. MAD measures the average absolute difference between the correctly predicted metal positions, 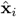, and the corresponding experimental positions, **x**_*i*_, and is computed as:

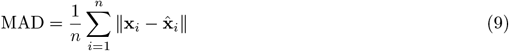

where *n* is the number of predicted metal sites. Lower MAD values indicate a closer match to the true positions, providing a complementary measure to precision and recall in assessing the overall performance of the method. These metrics enable a comprehensive comparison between SuperMetal and other existing methods.

## Data and Code Availability

The source code for SuperMetal is available on GitHub (https://github.com/XiaoboLinlin/SuperMetal/).

## Supporting Information

Supplementary information is available.

## Acknowledgements

Z.S. thanks the support of the Vanderbilt Data Science Postdoctoral Fellowship. X.L. and X.K. are grateful for the research funding and support provided by the Vanderbilt Data Science Institute. X.L. and P.T.C. thank the John R. Hall Professorship Endowment in Chemical Engineering for its support. J.L. expresses gratitude for the project opportunity provided by the Vanderbilt Data Science Institute. We sincerely thank Umang Chaudhry for facilitating access to these resources. We also acknowledge the computational resources (DGX A100) provided by the Vanderbilt Data Science Institute. J.M. is supported by a Humboldt Professorship of the Alexander von Humboldt Foundation. J.M. acknowledges funding by the Deutsche Forschungsgemeinschaft (DFG) through SFB1423 (421152132), SFB 1664 (514901783), TRR (514664767), and SPP 2363 (460865652). J.M. is supported by the Federal Ministry of Education and Research (BMBF) through the Center for Scalable Data Analytics and Artificial Intelligence (ScaDS.AI), through the German Network for Bioinformatics Infrastructure (de.NBI), and through the German Academic Exchange Service (DAAD) via the School of Embedded Composite AI (SECAI 15766814). J.M.’s Work in the Meiler laboratory is further supported through the National Institute of Health (NIH) through R01 HL122010, R01 DA046138, R01 AG068623, U01 AI150739, R01 CA227833, R01 LM013434, S10 OD016216, S10 OD020154, S10 OD032234.

## Author contributions

Z.S. conceptualized and designed the study and prepared the dataset. X.L., Z.S., and J.L. contributed to software development, model training, and analysis. X.L., Z.S., Y.L., J.L., X.K., J.S., J.M., and P.T.C. contributed to the writing and review of the manuscript. J.S., J.M., and P.T.C. provided funding support.

## References

1. Nanjiang Shu, Tuping Zhou, and Sven Hovmöller. Prediction of zinc-binding sites in proteins from sequence. Bioinformatics, 24(6):775–782, 2008.

2. Claudia Andreini, Lucia Banci, Ivano Bertini, and Antonio Rosato. Counting the zinc-proteins encoded in the human genome. Journal of proteome research, 5(1):196–201, 2006.

3. Keith A McCall, Chih-chin Huang, and Carol A Fierke. Function and mechanism of zinc metalloenzymes. The Journal of nutrition, 130(5):1437S–1446S, 2000.

4. Jeremy M Berg and Yigong Shi. The galvanization of biology: a growing appreciation for the roles of zinc. Science, 271(5252):1081–1085, 1996.

5. Maria Inês Costa, Ana Bela Sarmento-Ribeiro, and Ana Cristina Gonçalves. Zinc: from biological functions to therapeutic potential. International Journal of Molecular Sciences, 24(5):4822, 2023.

6. Christos T Chasapis, Ariadni C Loutsidou, Chara A Spiliopoulou, and Maria E Stefanidou. Zinc and human health: an update. Archives of toxicology, 86:521–534, 2012.

7. Maarten L Hekkelman, Ida de Vries, Robbie P Joosten, and Anastassis Perrakis. Alphafill: enriching alphafold models with ligands and cofactors. Nature Methods, 20(2):205–213, 2023.

8. Yu-Feng Lin, Chih-Wen Cheng, Chung-Shiuan Shih, Jenn-Kang Hwang, Chin-Sheng Yu, and Chih-Hao Lu. Mib: metal ion-binding site prediction and docking server. Journal of chemical information and modeling, 56(12):2287–2291, 2016.

9. Aditi Shenoy, Yogesh Kalakoti, Durai Sundar, and Arne Elofsson. M-ionic: prediction of metal-ion-binding sites from sequence using residue embeddings. Bioinformatics, 40(1):btad782, 2024.

10. José-Emilio Sánchez-Aparicio, Laura Tiessler-Sala, Lorea Velasco-Carneros, Lorena Roldán-Martín, Giuseppe Sciortino, and Jean-Didier Maréchal. Biometall: identifying metal-binding sites in proteins from backbone preorganization. Journal of Chemical Information and Modeling, 61(1):311–323, 2020.

11. Simon L Dürr, Andrea Levy, and Ursula Rothlisberger. Metal3d: a general deep learning framework for accurate metal ion location prediction in proteins. Nature Communications, 14(1):2713, 2023.

12. Elizabeth Brunk and Ursula Rothlisberger. Mixed quantum mechanical/molecular mechanical molecular dynamics simulations of biological systems in ground and electronically excited states. Chemical reviews, 115(12):6217–6263, 2015.

13. Zongfan Yang, Rebecca M Twidale, Silvia Gervasoni, Reynier Suardíaz, Charlotte K Colenso, Eric JM Lang, James Spencer, and Adrian J Mulholland. Multiscale workflow for modeling ligand complexes of zinc metalloproteins. Journal of Chemical Information and Modeling, 61(11):5658–5672, 2021.

14. Sharon L Guffy, Bryan S Der, and Brian Kuhlman. Probing the minimal determinants of zinc binding with computational protein design. Protein Engineering, Design and Selection, 29(8):327–338, 2016.

15. Bryan S Der, Mischa Machius, Michael J Miley, Jeffrey L Mills, Thomas Szyperski, and Brian Kuhlman. Metal-mediated affinity and orientation specificity in a computationally designed protein homodimer. Journal of the American Chemical Society, 134(1):375–385, 2012.

16. Lin Frank Song, Arkajyoti Sengupta, and Kenneth M Merz Jr. Thermodynamics of transition metal ion binding to proteins. Journal of the American Chemical Society, 142(13):6365–6374, 2020.

17. Haiyan Liu, Marcus Elstner, Efthimios Kaxiras, Thomas Frauenheim, Jan Hermans, and Weitao Yang. Quantum mechanics simulation of protein dynamics on long timescale. Proteins: Structure, Function, and Bioinformatics, 44(4):484–489, 2001.

18. Florent Barbault and Francois Maurel. Simulation with quantum mechanics/molecular mechanics for drug discovery. Expert Opinion on Drug Discovery, 10(10):1047–1057, 2015.

19. Andreas Zamanos, George Ioannakis, and Ioannis Z Emiris. Hydraprot: A new deep learning tool for fast and accurate prediction of water molecule positions for protein structures. Journal of Chemical Information and Modeling, 64(7):2594–2611, 2024.

20. Sangwoo Park and Chaok Seok. Galaxywater-cnn: Prediction of water positions on the protein structure by a 3d-convolutional neural network. Journal of Chemical Information and Modeling, 62(13):3157– 3168, 2022.

21. Ahmadreza Ghanbarpour, Amr H Mahmoud, and Markus A Lill. Instantaneous generation of protein hydration properties from static structures. Communications Chemistry, 3(1):188, 2020.

22. Kochi Sato, Mao Oide, and Masayoshi Nakasako. Prediction of hydrophilic and hydrophobic hydration structure of protein by neural network optimized using experimental data. Scientific reports, 13(1):2183, 2023.

23. Zhijian Liu, Haotian Tang, Yujun Lin, and Song Han. Point-voxel cnn for efficient 3d deep learning. Advances in neural information processing systems, 32, 2019.

24. Cheng Wang, Ming Cheng, Ferdous Sohel, Mohammed Bennamoun, and Jonathan Li. Normalnet: A voxel-based cnn for 3d object classification and retrieval. Neurocomputing, 323:139–147, 2019.

25. Yang Zhang, Wenbing Huang, Zhewei Wei, Ye Yuan, and Zhaohan Ding. Equipocket: an e (3)-equivariant geometric graph neural network for ligand binding site prediction. arXiv preprint 2302.12177, 2023.

26. Sangwoo Park. Water position prediction with se (3)-graph neural network. bioRxiv, pages 2024–03, 2024.

27. Yang Song, Jascha Sohl-Dickstein, Diederik P Kingma, Abhishek Kumar, Stefano Ermon, and Ben Poole. Score-based generative modeling through stochastic differential equations. arXiv preprint 2011.13456, 2020.

28. Gabriele Corso, Hannes Stärk, Bowen Jing, Regina Barzilay, and Tommi Jaakkola. Diffdock: Diffusion steps, twists, and turns for molecular docking. arXiv preprint 2210.01776, 2022.

29. Mohamed Amine Ketata, Cedrik Laue, Ruslan Mammadov, Hannes Stärk, Menghua Wu, Gabriele Corso, Céline Marquet, Regina Barzilay, and Tommi S Jaakkola. Diffdock-pp: Rigid protein-protein docking with diffusion models. arXiv preprint 2304.03889, 2023.

30. Joseph L Watson, David Juergens, Nathaniel R Bennett, Brian L Trippe, Jason Yim, Helen E Eisenach, Woody Ahern, Andrew J Borst, Robert J Ragotte, Lukas F Milles, et al. De novo design of protein structure and function with rfdiffusion. Nature, 620(7976):1089–1100, 2023.

31. Emiel Hoogeboom, Vıctor Garcia Satorras, Clément Vignac, and Max Welling. Equivariant diffusion for molecule generation in 3d. In International conference on machine learning, pages 8867–8887. PMLR, 2022.

32. Karsten Kreis, Tim Dockhorn, Zihao Li, and Ellen Zhong. Latent space diffusion models of cryo-em structures. arXiv preprint 2211.14169, 2022.

33. Dominik JE Waibel, Ernst Röell, Bastian Rieck, Raja Giryes, and Carsten Marr. A diffusion model predicts 3d shapes from 2d microscopy images. In 2023 IEEE 20th International Symposium on Biomedical Imaging (ISBI), pages 1–5. IEEE, 2023.

34. Zhiye Guo, Jian Liu, Yanli Wang, Mengrui Chen, Duolin Wang, Dong Xu, and Jianlin Cheng. Diffusion models in bioinformatics and computational biology. Nature reviews bioengineering, 2(2):136–154, 2024.

35. Geoffrey Roeder, Yuhuai Wu, and David K Duvenaud. Sticking the landing: Simple, lower-variance gradient estimators for variational inference. Advances in Neural Information Processing Systems, 30, 2017.

36. Luke Metz, Ben Poole, David Pfau, and Jascha Sohl-Dickstein. Unrolled generative adversarial networks. arXiv preprint 1611.02163, 2016.

37. Franco Scarselli, Marco Gori, Ah Chung Tsoi, Markus Hagenbuchner, and Gabriele Monfardini. The graph neural network model. IEEE transactions on neural networks, 20(1):61–80, 2008.

38. Zeming Lin, Halil Akin, Roshan Rao, Brian Hie, Zhongkai Zhu, Wenting Lu, Nikita Smetanin, Robert Verkuil, Ori Kabeli, Yaniv Shmueli, et al. Evolutionary-scale prediction of atomic-level protein structure with a language model. Science, 379(6637):1123–1130, 2023.

39. Martin Ester, Hans-Peter Kriegel, Jörg Sander, and Xiaowei Xu. Density-based spatial clustering of applications with noise. In Int. Conf. knowledge discovery and data mining, volume 240, 1996.

40. José Jiménez, Stefan Doerr, Gerard Martínez-Rosell, Alexander S Rose, and Gianni De Fabritiis. Deepsite: protein-binding site predictor using 3d-convolutional neural networks. Bioinformatics, 33(19):3036–3042, 2017.

41. Chih-Hao Lu, Chih-Chieh Chen, Chin-Sheng Yu, Yen-Yi Liu, Jia-Jun Liu, Sung-Tai Wei, and Yu-Feng Lin. Mib2: metal ion-binding site prediction and modeling server. Bioinformatics, 38(18):4428–4429, 2022.

42. Mario Geiger and Tess Smidt. e3nn: Euclidean neural networks. arXiv preprint 2207.09453, 2022.

43. Michael Buckland and Fredric Gey. The relationship between recall and precision. Journal of the American society for information science, 45(1):12–19, 1994.

44. Josh Abramson, Jonas Adler, Jack Dunger, Richard Evans, Tim Green, Alexander Pritzel, Olaf Ronneberger, Lindsay Willmore, Andrew J Ballard, Joshua Bambrick, et al. Accurate structure prediction of biomolecular interactions with alphafold 3. Nature, pages 1–3, 2024.

45. Amr Moustafa. Structural and Functional Analyses of Secretory and Excretory Proteins from Onchocerca volvulus as Basis for Rational Drug Design. PhD thesis, Staats-und Universitätsbibliothek Hamburg Carl von Ossietzky, 2016.

46. Lara S Kallander, David Washburn, Mark A Hilfiker, Hilary Schenck Eidam, Brian G Lawhorn, Joanne Prendergast, Ryan Fox, Sarah Dowdell, Sharada Manns, Tram Hoang, et al. Reverse hydroxamate inhibitors of bone morphogenetic protein 1. ACS Medicinal Chemistry Letters, 9(7):736–740, 2018.

47. Xiaohan Kuang, Zhaoqian Su, Yunchao Liu, Xiaobo Lin, Jesse Spencer Smith, Tyler Derr, Yinghao Wu, and Jens Meiler. Superwater: Predicting water molecule positions on protein structures by generative ai. bioRxiv, pages 2024–11, 2024.

48. Yannis Nevers, Natasha M Glover, Christophe Dessimoz, and Odile Lecompte. Protein length distribution is remarkably uniform across the tree of life. Genome Biology, 24(1):135, 2023.

49. Sam M Ireland and Andrew CR Martin. Zincbind—the database of zinc binding sites. Database, 2019:baz006, 2019.

50. Peter W Rose, Andreas Prlić, Ali Altunkaya, Chunxiao Bi, Anthony R Bradley, Cole H Christie, Luigi Di Costanzo, Jose M Duarte, Shuchismita Dutta, Zukang Feng, et al. The rcsb protein data bank: integrative view of protein, gene and 3d structural information. Nucleic acids research, page gkw1000, 2016.

51. Geng-Yu Lin, Yu-Cheng Su, Yen Lin Huang, and Kun-Yi Hsin. Mespeus: a database of metal coordination groups in proteins. Nucleic Acids Research, 52(D1):D483–D493, 2024.

52. Jonathan Ho, Ajay Jain, and Pieter Abbeel. Denoising diffusion probabilistic models. Advances in neural information processing systems, 33:6840–6851, 2020.

53. Ilia Igashov, Hannes Stärk, Clément Vignac, Arne Schneuing, Victor Garcia Satorras, Pascal Frossard, Max Welling, Michael Bronstein, and Bruno Correia. Equivariant 3d-conditional diffusion model for molecular linker design. Nature Machine Intelligence, pages 1–11, 2024.

54. Yang Song and Stefano Ermon. Generative modeling by estimating gradients of the data distribution. Advances in neural information processing systems, 32, 2019.

55. Yang Song and Stefano Ermon. Improved techniques for training score-based generative models. Advances in neural information processing systems, 33:12438–12448, 2020.

56. Alexander Quinn Nichol and Prafulla Dhariwal. Improved denoising diffusion probabilistic models. In International conference on machine learning, pages 8162–8171. PMLR, 2021.

57. FM Bi, WK Wang, and L Chen. Dbscan: density-based spatial clustering of applications with noise. J. Nanjing Univ, 48(4):491–498, 2012.

58. Fabian Pedregosa, Gaël Varoquaux, Alexandre Gramfort, Vincent Michel, Bertrand Thirion, Olivier Grisel, Mathieu Blondel, Peter Prettenhofer, Ron Weiss, Vincent Dubourg, et al. Scikit-learn: Machine learning in python. the Journal of machine Learning research, 12:2825–2830, 2011.

