## supplemental information for "SuperMetal: A Generative AI Framework for Rapid and Precise Metal Ion Location Prediction in Proteins"

### 1 SuperMetal Architecture

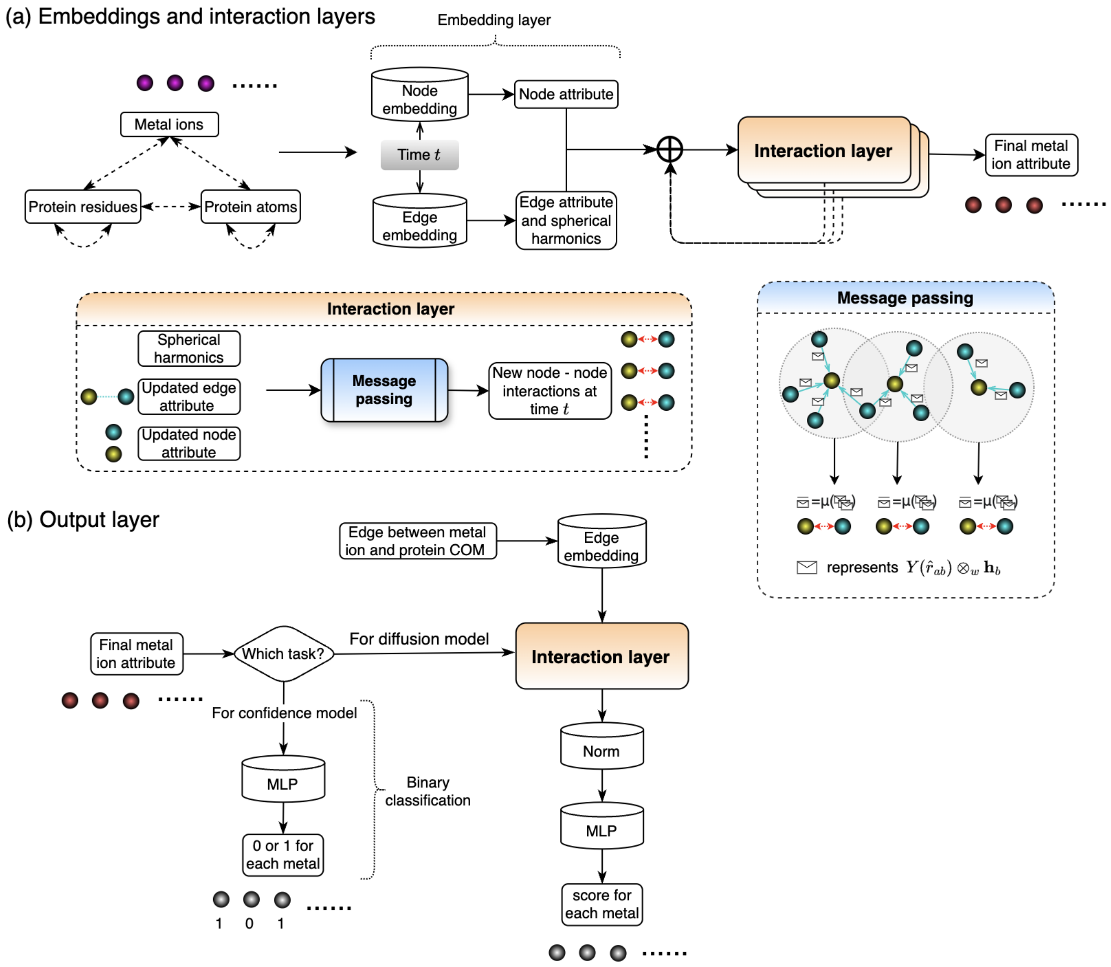


**Fig. S1** **Overview of the SuperMetal Model Architecture.** **(a) Embedding and interaction layers:** This section shows the input metal ions, protein residues, and atoms, where the message-passing module features the center node $a$ (in yellow) and its surrounding nodes $b$ (in blue). Node and edge embeddings, along with spherical harmonics, are incorporated to update node interactions at time $t$. The operation $\otimes_w$ represents the spherical tensor product of irreducible representations of $SO(3)$, weighted by path coefficients $w$. **(b) Output layers:** Metal ion attributes from the embedding and interaction layers are used to compute scores for the diffusion model or generate binary classification labels for the confidence model.

Our SuperMetal architecture is adapted from DiffDock[1], with key modifications tailored for metal-binding site prediction. The score model $s(\mathbf{x}, \mathbf{y}, t)$ and confidence model $d(\mathbf{x}, \mathbf{y})$ take as input the current metal ion positions $\mathbf{x}$ and the protein structure $\mathbf{y}$ in 3D space. Both models output SE(3)-invariant vectors, ensuring that the predicted metal ion positions are relative to the protein structure, which may be located arbitrarily. The output of the score model is located in the tangent space $T_r T_3$, equivalent to $\mathbb{R}^3$, and produces SE(3)-equivariant translational score predictions. Both the score and confidence models are based on SE(3)-equivariant convolutional networks, adapted from prior works such as [2, 3], but designed specifically for metal-binding predictions. Both models utilize detailed all-atom protein structures and operate on a multiscale representation, including a coarse-grained representation using $\alpha$-carbon atoms for enhanced performance. Moreover, the SE(3)-equivariant architecture consistently handles rotations and translations, enabling the model to capture relative geometric relationships instead of relying on a fixed orientation or alignment. By leveraging the underlying physics and geometry—rather than memorizing explicit motifs—this approach mitigates overfitting to repetitive patterns and enhances the model's ability to generalize to novel protein environments.

As illustrated in Fig. S1, the architecture begins with the embedding layer, where protein and metal structures are represented as heterogeneous geometric graphs. Nodes in these graphs represent metal ions, protein residues (centered on the $\alpha$-carbon), and protein atoms. These nodes are connected using sparse distance cutoffs that vary based on diffusion time. The radii used for these cutoffs are derived from [1]. Node interactions include receptor atoms with other receptor atoms, receptor atoms with metal ions, receptor residues with other receptor residues, and receptor residues with metal ions, while interactions between metal ions are excluded due to their generally larger distances. The nodes are initialized with features that include categorical information about the protein residues and atoms, enhanced with embeddings derived from the protein sequence language model, ESMFold [4]. These node features are then augmented with sinusoidal embeddings representing diffusion time and processed by multilayer perceptrons (MLPs) to generate scalar features that are passed into the interaction layers. Similarly, edge features, encoding the distances between nodes, are embedded using a Gaussian smear and concatenated with sinusoidal diffusion time embeddings, then processed via MLPs. Spherical harmonic representations of edge vectors are calculated to capture geometric relationships between nodes.

In the interaction layer, as previously discussed, cutoff distances are established for each type of node, allowing each node to receive messages from the tensor product of its nearby connected nodes and the spherical harmonic representations of the corresponding edge vectors. The weights for this tensor product are computed using MLPs, based on the edge embeddings and the scalar features of the nodes. Consequently, each node aggregates messages from its neighboring nodes and edges, and these messages are averaged to update the interactions for the ”leading” node. For a more detailed explanation, please refer to ref. [1]. As illustrated in Fig. S1, after several iterations of interaction layer processing, the various interactions are updated, leading to refined node features, including those for metal ions. These updated metal ion features are then utilized in the final layer to compute a score for each metal ion.

In the confidence model, the final layer produces a scalar output representing the confidence for each metal ion through a fully connected layer. For the score model, translational scores corresponding to the linear acceleration of the metal ions are computed. Each metal ion is convolved with the (unweighted) center of mass $c$ of the protein. The process is described as follows:

$$\begin{aligned} \mathbf{v} &\leftarrow Y(\hat{r}_{ca}) \otimes_{\psi_{ca}} \mathbf{h}_a \\ \text{with} \quad \psi_{ca} &= \Psi(\mu(r_{ca}), \mathbf{h}_a^0) \end{aligned} \tag{1}$$

Here, $\mathbf{v}$ represents the output vector, $Y$ corresponds to spherical harmonics up to $\ell = 2$, and $\mathbf{h}_a$ is the node embedding of node $a$. The operation $\otimes_{\psi_{ca}}$ denotes the spherical tensor product of irreducible representations with path weights $\psi_{ca}$, computed by an MLP $\Psi$. $\mu(r_{ca})$ represents radial basis embeddings of the edge length, and $\mathbf{h}_a^0$ refers to the initial scalar features of node $a$. After obtaining $\mathbf{v}$, the vector is further refined using an MLP that takes the magnitude of the vector and sinusoidal embeddings of the diffusion time as input. Finally, the outputs are multiplied by $1/\sigma_{tr}$ to recover the translational scores.

#### 2 SuperMetal Inference and Prediction

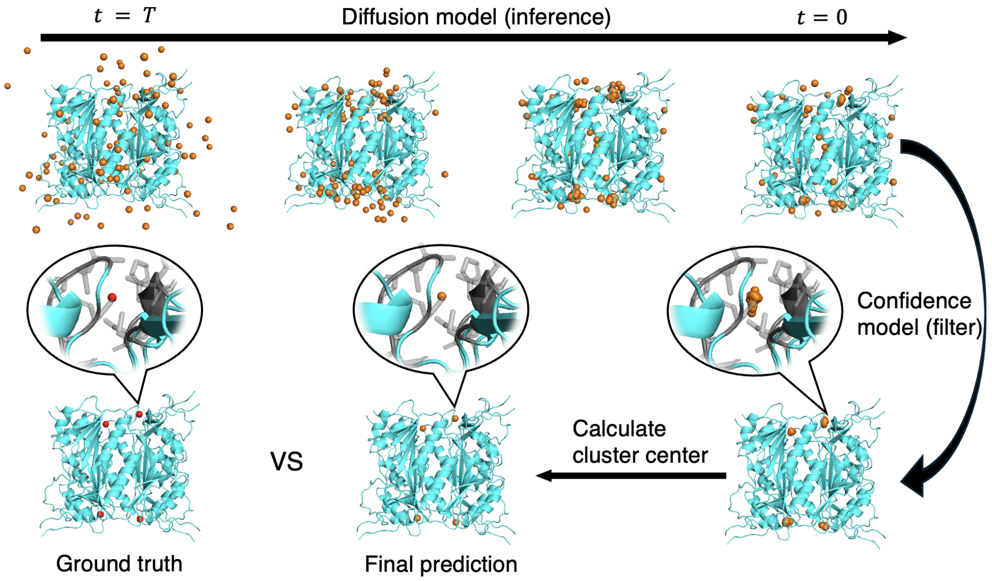


**Fig. S2 Overview of the metal ion prediction process in SuperMetal.** The process begins by randomly positioning the metal ions based on a normal distribution at time $t = T$. As the system evolves towards $t = 0$, the metal ions progressively move towards biologically relevant locations. A confidence model is applied to filter out low-confidence or erroneous predictions. The final metal ion positions are determined by calculating the cluster centers of the high-confidence predictions.

During the inference step, 100 metal ions are randomly distributed across the system at time $t = T$, and their positions are refined through the application of reverse diffusion, which operates over translational degrees of freedom. The positions generated are subsequently refined using a confidence model trained to select the most probable locations for the metal ions. These refined predictions tend to cluster around actual metal-binding sites, as illustrated in Fig. S2. The final predicted positions are determined by computing the centroid of each cluster, ensuring accurate placement of the metal ions in the protein structure.

### References


[1] Gabriele Corso, Hannes Stärk, Bowen Jing, Regina Barzilay, and Tommi Jaakkola. Diffdock: Diffusion steps, twists, and turns for molecular docking. *arXiv preprint arXiv:2210.01776*, 2022.

[2] Nathaniel Thomas, Tess Smidt, Steven Kearnes, Lusann Yang, Li Li, Kai Kohlhoff, and Patrick Riley. Tensor field networks: Rotation-and translation-equivariant neural networks for 3d point clouds. *arXiv preprint arXiv:1802.08219*, 2018.

[3] Mario Geiger and Tess Smidt. e3nn: Euclidean neural networks. *arXiv preprint arXiv:2207.09453*, 2022.

[4] Zeming Lin, Halil Akin, Roshan Rao, Brian Hie, Zhongkai Zhu, Wenting Lu, Nikita Smetanin, Robert Verkuil, Ori Kabeli, Yaniv Shmueli, et al. Evolutionary-scale prediction of atomic-level protein structure with a language model. *Science*, 379(6637):1123–1130, 2023.
